# Differential stimulation of pluripotent stem cell-derived human microglia leads to exosomal proteomic changes affecting neurons

**DOI:** 10.1101/2021.07.19.452610

**Authors:** Anna Mallach, Johan Gobom, Charles Arber, Thomas M Piers, John Hardy, Selina Wray, Henrik Zetterberg, Jennifer Pocock

## Abstract

Microglial exosomes are an emerging communication pathway, implicated in fulfilling homeostatic microglial functions and transmitting neurodegenerative signals. Gene variants of the *triggering receptor expressed myeloid cells-2 (TREM2)* are associated with an increased risk of developing dementia. We investigated the influence of the *TREM2* Alzheimer’s disease risk variant, R47H^het^, on the microglial exosomal proteome consisting of 3019 proteins secreted from human iPS-derived microglia (iPS-Mg). Exosomal protein content changed according to how iPS-Mg were stimulated. Thus lipopolysaccharide (LPS) induced microglial exosomes to contain more inflammatory signals, whilst stimulation with the TREM2 ligand phosphatidylserine (PS^+^) increased metabolic signals within the microglial exosomes. We tested the effect of these exosomes on neurons and found that the exosomal protein changes were functionally relevant and influenced downstream functions in both neurons and microglia. Exosomes from R47H^het^ iPS-Mg contained disease-associated microglial (DAM) signature proteins, and were less able to promote outgrowth of neuronal processes and increase mitochondrial metabolism in neurons compared with exosomes from *TREM2* common variant iPS-Mg. Taken together these data highlight the importance of microglial exosomes in fulfilling microglial functions. Additionally, variations in the exosomal proteome influenced by the R47H^het^ *TREM2* variant may underlie the increased risk of Alzheimer’s disease associated with this variant.

**Graphical Abstract:** 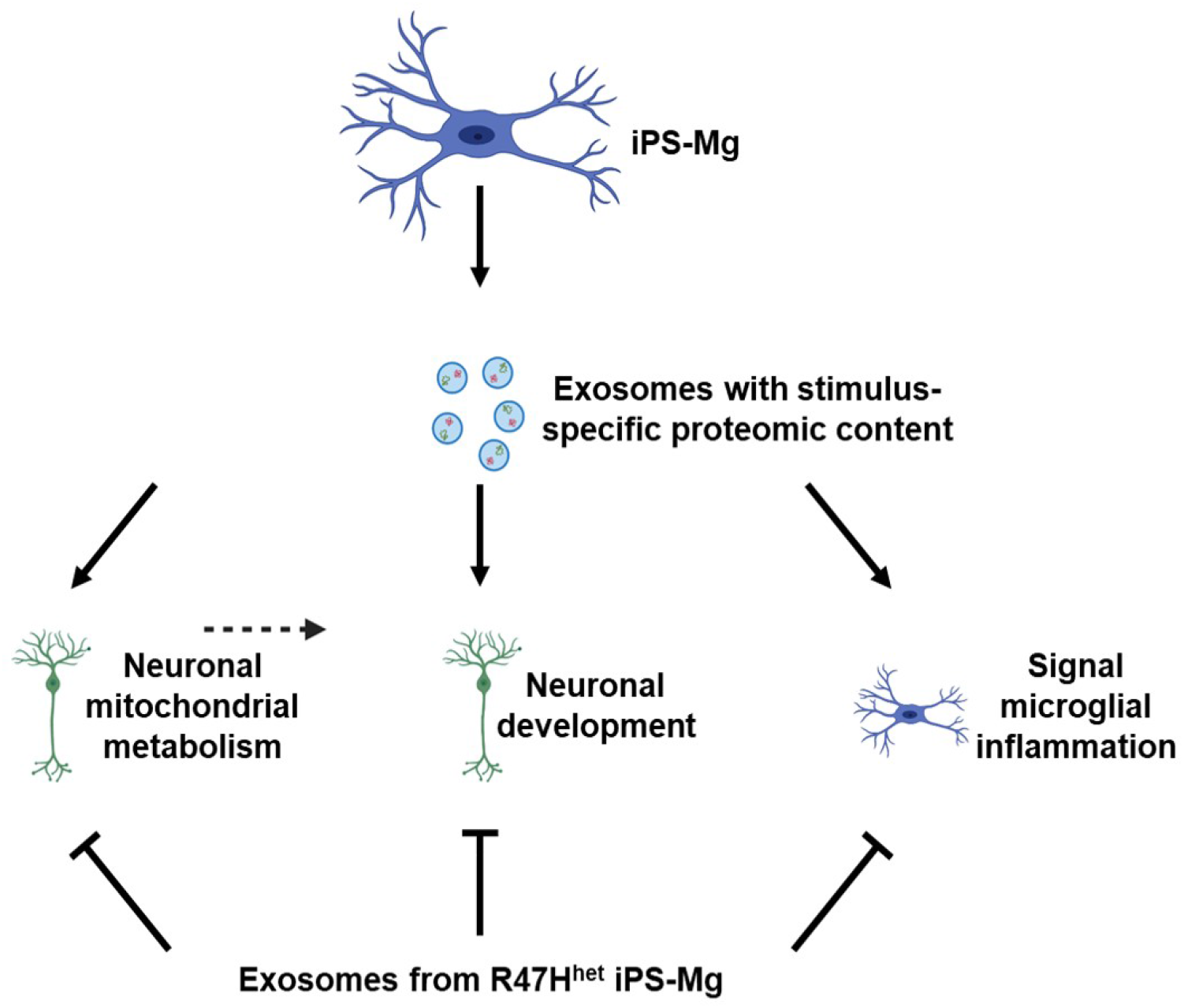

## Introduction

Recent studies have identified late-onset Alzheimer’s disease (LO-AD) risk genes associated with microglia, the immune cells of the brain [1–5]. One of these genes encodes the protein, triggering receptor expressed on myeloid cells-2 (TREM2), which is expressed on microglia in the CNS and senses lipids, such as phospholipids, in the extracellular environment [6–8]. The R47H^het^ *TREM2* variant significantly increases the risk of developing LO-AD [3,9]. TREM2 deficits have been linked to reduced phagocytosis, metabolism and an aberrant response to extracellular stimuli [10–13], impairing key microglial functions.

One key microglial function is the secretion of a range of factors, including exosomes [14,15]. Exosomes are small extracellular vesicles, which can be secreted from different cells, including microglia. Microglial exosomes have been implicated in the progression of neurodegeneration [16–19] and neuronal functioning including neurite outgrowth [20,21]. Exosomal content has also been shown to change depending on the signals microglia receive [15].

We have recently shown that the R47H^het^ *TREM2* variant decreased the secretion of exosomes from patient-derived induced pluripotent stem cell-derived microglia (iPS-Mg) [22]. The R47H^het^ *TREM2* variant also affected the ability of exosomes to rescue neurons from cell death compared with exosomes secreted from common variant (Cv) *TREM2* expressing iPS-Mg [22].

In this study, we investigated the influence of TREM2 on exosomal content using patient iPS-Mg, expressing Cv or R47H^het^ *TREM2*, by assessing network changes of 3019 proteins rather than the analysis of top most abundant proteins as was determined in our previous paper [22]. Thus it was revealed that LPS treatment of iPS-Mg induced a specific inflammatory response in exosomes, whilst PS^+^ induced changes in metabolic exosomal proteins. Exosomes from R47H^het^ iPS-Mg added to neurons were less capable of supporting neuronal development and metabolic pathways compared with Cv exosomes, whilst both Cv and R47H^het^ exosomes were able to transmit inflammatory information to other homeostatic microglia in an autocrine signalling pathway.

## Materials and Methods

### Cell lines

R47H^het^ fibroblasts from two different patients were obtained through a material transfer agreement with the University of California Irvine Alzheimer’s Disease Research Centre. iPS lines were generated as previously described [11]. For this study, three clones per patient line were used, of R47H^het^, 6 cell lines in total. In addition the following *TREM2* common variant lines were used to generate iPS-Mg; CTRL1 (kindly provided by Prof S Wray, UCL Queen Square Institute of Neurology), CTRL2 (SBAD03, StemBANCC), CTRL3 (SFC840, StemBANCC) and CTRL4 (BIONi010-C, EBiSC). To generate iPS-neurons, the iPS line RBi001-a (Sigma Aldrich) was used.

### Cell culture

#### iPS-Mg

Using our previously published protocol and iPSC lines, iPS-Mg were generated [11,23]. Briefly, embryoid bodies were generated from iPSC using IL-3, MCSF and β-mercaptoethanol using a previously described protocol [10,24]. Myeloid progenitors were selected and further differentiated using IL-34, MCSF and TGF-β for two weeks, in addition to CX3CL1 and CD200 for the final 3 days [11,23]. The generated iPS-Mg have previously been characterised in terms of their microglial signature gene expression, and display typical microglial functions, such as phagocytosis of particles, intracellular signalling, and responses to inflammatory stimuli [11,25].

#### iPS-neurons

Differentiation of iPSC into iPS-neurons was performed following a published protocol [26]. In brief, confluent iPSC were switched to N2B27 media, supplemented with 10 µM SB431542 (Tocris) and 1 µM dorsomorphin (Tocris). N2B27 medium was composed of 1:1 DMEM-F12 and Neurobasal medium, containing 0.5x N2 supplement, 0.5x B27 supplement, 0.5x NEAA, 1 mM L-glutamine, 25 U pen/strep, 10 µM β-mercaptoethanol and 25 U insulin. After 10 days of neural induction, the cells were maintained in N2B27 without SB431542 and dorsomorphin until after day 100, at which time the experiments were performed. The differentiation of the iPS-neurons have been previously characterised [27].

#### SH-SY5Y neurons

SH-SY5Y cells (a kind gift from Prof R de Silva, UCL Queen Square Institute of Neurology), were differentiated using a previously established protocol [28] by incubation with retinoic acid (RA, 10 µM) for 5 days and brain derived neurotrophic factor (BDNF, 50 nM) for 7 days. These cells were differentiated using a previously established protocol [28] by incubation with retinoic acid (RA, 10 µM) for 5 days and BDNF (50 nM) for 7 days. The differentiated cells are an established neuron-like model displaying a metabolic profile similar to primary neurons and referred to as SH-SY5Y neurons in this paper [29].

### Cell treatment and exosome collection

PS^+^ cells were generated through subjecting SH-SY5Y to a heat shock for 2 h at 45°C [10]. The heat-shocked neurons express phosphatidylserine on their outer plasma membrane, which we have verified previously verified [10,25], a known ligand of TREM2 [8,30]. Therefore, when PS^+^ neurons were added to iPS-Mg, they were termed PS^**+**^.

iPS-Mg were treated with lipopolysaccharide (LPS, 100 ng/ml) or 2:1 PS^+^:iPS-Mg for 24 h before the supernatant was collected. Exosomes were extracted from supernatant with an ExoQuick kit (System Biosciences). The presence of exosomal markers and size distribution of the particles was confirmed previously [22]. Extracted exosomes were lysed in RIPA buffer and their protein content determined by BCA analysis.

SH-SY5Y neurons, iPS-Mg or iPS-neurons were exposed to exosomes from Cv or R47H variant iPS-Mg. For this, 6 µg of exosomal protein were added to iPS-Mg, SH-SY5Y neurons or iPS-neurons for 24 h. For iPS-Mg experiments, exosomes from the same genetic background were added, in that Cv iPS-Mg were treated with Cv exosomes and R47H^het^ iPS-Mg were treated with R47H^het^ exosomes.

### Proteomic analysis

Proteomic data from 50 µg of iPS-Mg exosomes for each treatment was generated from liquid chromatography tandem mass spectrometry (LC-MS/MS) experiments described previously [22] were further analysed to elucidate changes in exosomal protein content depending on cell treatment and *TREM2* variant. Different repeats were pooled into three independent samples. Functional analysis was performed using FunRich software. Heatmaps and principal component analysis (PCA) plots were produced with MATLAB.

Based on previously published weighted gene co-expression network analysis [31,32], two signed networks were constructed, one for each of the genotypes, using different treatments as traits. The code, run through RStudio, was based on the tutorial provided with the R package. Briefly, proteins were clustered based on their dissimilarity measure before being raised to a power function, based on the assumption of a scale free network. For both networks, this power was 30. To merge closely related modules, 30 proteins were set as a minimum module size and a threshold of 0.25 chosen to merge closely clustered modules. Module preservation was performed using previously published papers [33], using *medianRank, Zsummary* and kME correlations [33,34].

Functional annotation of the modules was performed with the “GOenrichmentAnalysis” package, provided through RStudio, with a Bonferroni correction applied to the ρ value to account for multiple comparisons.

### Quantitative PCR

Quantitative PCR (qPCR) experiments were performed to determine the effect of exosomes on iPS-Mg and neurons. Cell treated with exosomes as above were lysed in Trizol. RNA was extracted using the DirectZol RNA MiniPrep Plus kit (Cambridge Bioscience, #74106). cDNA was generated with a high-capacity cDNA reverse transcription kit (Applied Biosystems, #4368814) and qPCR analysis performed using Taqman Universal Master Mix (Life Technologies, #4440038) using specific primers (Table 1) in an Eppendorf Mastercycler.

**Table 1:** Probes used for qPCR.

| Gene name | Identifier |
| --- | --- |
| **APOE** | Hs00171168_m1 |
| **AXL** | Hs01064444_m1 |
| **CSF1R** | Hs00911250_m1 |
| **GAPDH** | Hs02758991_g1 |
| **GPNUMB** | Hs01095669_m1 |
| **GRN** | Hs00963707_g1 |
| **HSPD1** | Hs01036753_g1 |
| **IL1B** | Hs00174097_m1 |
| **IL6** | Hs00985639_m1 |
| **PPARGC** | Hs01016719_m1 |
| **TNF** | Hs01113624_g1 |
| **TREM2** | Hs00219132_m1 |
| **TYROBP** | Hs00182426_m1 |

### Immunocytochemistry

iPS-neurons were exposed to basal exosomes, derived from untreated iPS-Mg, carrying either the Cv or R47H^het^ *TREM2* variant as described above and fixed with 4% PFA with 4% sucrose for 20 min. The cells were then quenched in 50 mM NH4Cl for 10 min, followed by permeabilisation with 0.2% Triton X-100 for 5 min. Primary antibodies, Tuj1 (BioLegend, #801202) and GAP43 (Abcam, #75810), were incubated overnight at 4°C in 5% normal goat serum. Appropriate secondary antibodies were used for 1 h at room temperature before the coverslips were mounted on slides using Vectashield with DAPI. Images were acquired on a Zeiss LSM710 confocal microscope using the LSM Pascal 5.0 software. Eight regions of interest (ROI) were taken per coverslip. For neuronal process length analysis, Tuj1 staining was skeletonised using an adapted version of a previously described protocol [35]. GAP43 levels were quantified using a Fiji macro based on a previously published study [36]. Briefly, the overlay between the GAP43 and Tuj1 images was calculated with the area of GAP43 staining within Tuj1 positive pixels normalised to total area of Tuj1 positive pixels plotted.

### Metabolic assays

Changes in metabolic activity were measured in SH-SY5Y following 24 h incubation with exosomes. Neurons were incubated with 0.5 mg/ml MTT solution for 2 h at 37°C. Afterwards the MTT solution was removed and the solvent, isopropanol, added for 15 min at room temperature to dissolve the MTT crystals. The plate was then read on a Tecan 10M plate reader. To normalise the results to total cell number, taking into account potential changes in cell viability, 50 µl crystal violet (Pro-Lab Diagnostics, #PL7000) were added to the plate, and excess removed after 20 min. After the plate was air-dried, methanol was added and the plate was read on a Tecan 10M plate reader again.

ATP levels were measured in neurons treated with exosomes for 24 h using a commercially available ATP kit (Thermo Fisher Scientific, #A22066). The cells were lysed in cell lysis reagent and centrifuged at 1,000g for 1 min. 10 µl of sample or ATP standard was added to 100 µl ATP reaction mix, after the background luminescence of the ATP reaction mix was read. Luminescence was read on a Tecan 10M. Data were normalised to the protein concentration of each sample, as determined through BCA assay.

To ensure that the changes measured in both assays were due to changes in metabolic activity instead of cell proliferation or cell death, the crystal violet data obtained following the MTT assay were plotted (**Supplementary Fig 7Bi**).

### Endotoxin assay

To test whether LPS was carried over in exosomal extractions from iPS-Mg exposed to LPS, endotoxin levels in exosomes and iPS-Mg medium were measured using the HEK Blue LPS detection kit 2 (InvivoGen) following manufacturers’ instructions. Briefly, 25,000 HEK Blue cells were exposed to different quantities of exosomes, medium or known levels of endotoxin. The release of secreted embryonic alkaline phosphatase from these cells following activation of NF-κB was determined through QUANTI-Blue and the plate was read on a Tecan 10M to measure absorbance.

### Statistical analysis

The results were represented as a mean of at least three separate experimental repeats ± SEM with a ρ value of 0.05 or below considered statistical significant. The results were analysed using Prism Software version 5, MATLAB or R Studio. Analysis was performed on pooled control lines and pooled R47H^het^ cell lines. As described previously [22], proteomic data were log10 transformed prior to analysis.

## Results

### Stimulatory treatments induce specific exosomal proteomic changes

Changes in exosomal proteins following treatment of microglia with have been reported previously [14,15]. In our previous study, we reported the top 200 most abundant and significantly changed exosomal proteins secreted from iPS-Mg [22], however we did not investigate subtle changes induced by the R47H^het^ *TREM2* variant and the effects of different iPS-Mg stimuli on the exosomal proteome.

The overall effect of different conditions on the exosomal proteome was determined by generating a heatmap that clustered the different samples based on exosomal protein levels (**Fig 1A**). Therefore, samples which were similar in terms of exosomal protein levels clustered closely together, allowing for a visual inspection of overall differences between the different treatment conditions. This showed that following treatment with LPS or PS^+^, exosomes formed treatment-specific clusters, independent of *TREM2* status. Furthermore, exosomes from basal (unstimulated) iPS-Mg were different between the Cv and R47H^het^ genotypes. These differences were shown when the proposed function of the proteins was analysed (**Fig 1B**). Whilst exosomes from basal Cv iPS-Mg contained higher levels of proteins involved in cell growth and/or maintenance, R47H^het^ basal exosomes contained more transcription-related proteins. This is consistent with our previous finding that the most abundant proteins in exosomes from R47H^het^ iPS-Mg were involved in the negative regulation of transcription [22]. Stimulus-specific changes were associated with increases in the cytoskeletal organization and biogenesis, and protein folding for LPS and PS^+^ respectively (**Fig 1B**) in both Cv and R47H^het^ derived exosomes.

**Figure 1.**
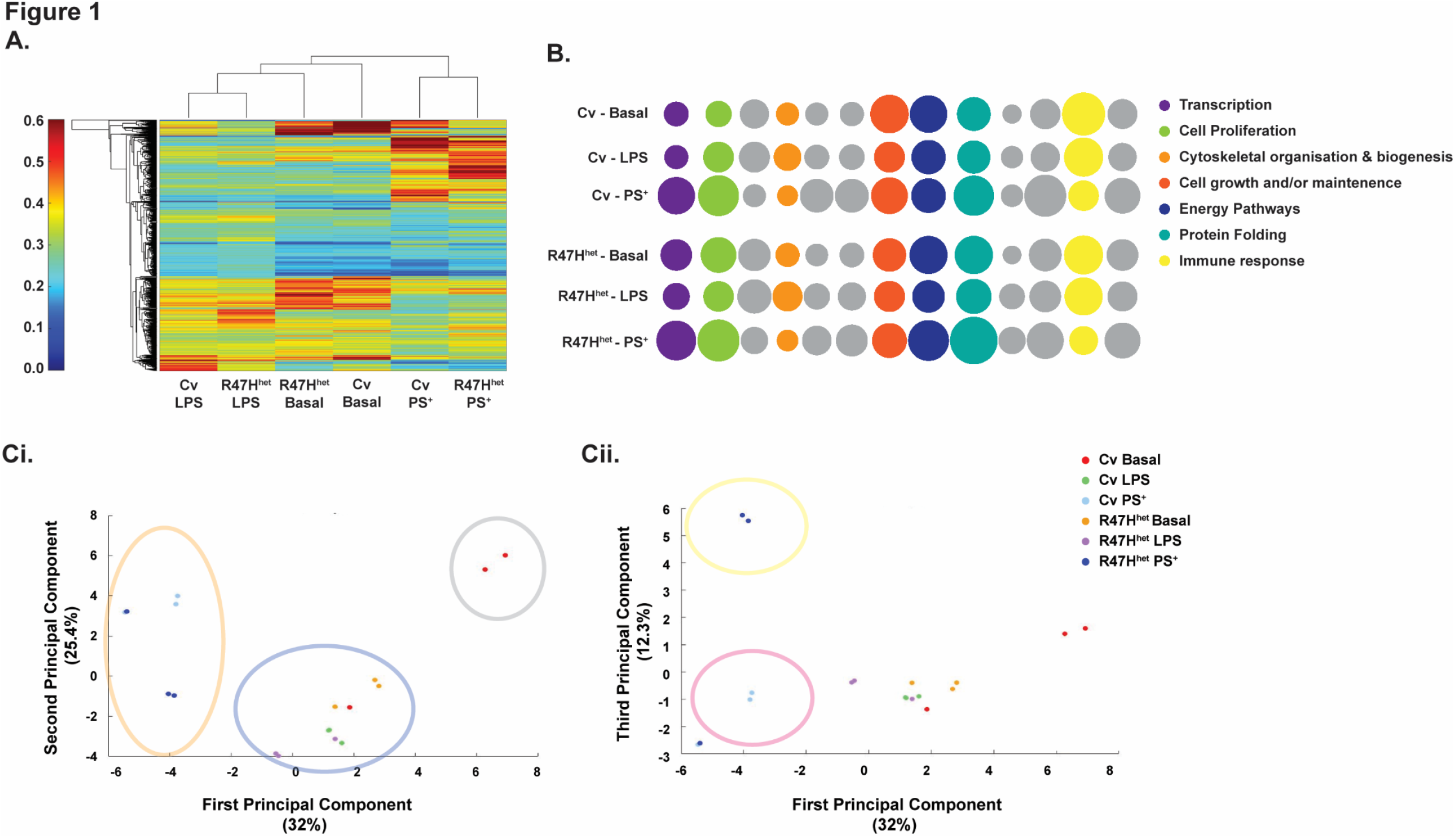
Following treatments, exosomes fall into specific clusters. Following LC-MS/MS process, a heatmap of exosomal proteins was constructed to analyse the effect of the different iPS-Mg treatments on exosomal proteome content (**A**). The relative abundance of proteins associated with different biological processes, identified with FunRich, was plotted for the different conditions with the diameter of the circles representing protein abundance (**B**). Principal component analysis (PCA) to seek clusters between the different samples was generated (**C**). N=3 independent samples analysed through LC-MS/MS for each condition.

To reduce the variance in the data, principal component analysis (PCA) was applied to further analyse the clusters observed in the heatmap (**Fig 1C**). In line with the heatmap (**Fig 1A**), the samples fell into separate LPS and PS^+^ clusters (**Fig 1Ci**, blue and red respectively). The basal Cv samples formed their own cluster (**Fig 1Ci**, grey), whilst the basal R47H^het^ samples fell into the LPS cluster. This confirmed the indication from the heatmap, that the basal R47H^het^ exosomes resemble exosomes from LPS-treated iPS-Mg. In an attempt to identify further clusters, the third principal component (PC) was also plotted (**Fig 1Cii**), also showing differences between the Cv and R47H^het^ exosomes extracted from iPS-Mg treated with PS^+^ in addition to the cluster of untreated Cv exosomes and exosomes from LPS-treated iPS-Mg.

Overall, exosomal content appeared to be significantly influenced by iPS-Mg treatment with these two different treatments inducing different proteomic changes. Furthermore, the *TREM2* R47H^het^ variant had an effect on exosomal content from untreated iPS-Mg and iPS-Mg treated with PS^+^.

### R47H^het^ exosomes contain more DAM-associated proteins

In addition to analysing the trends in all of the identified exosomal proteins, we investigated a subset of proteins identified in the exosomes. Due to the inverse relationship between TREM2 expression and the development of a disease-associated microglial (DAM) signature [37], we specifically probed the exosome dataset for DAM-related proteins, based on previous publications [37–39]. A number of DAM-associated proteins were found in exosomes, which were plotted in a heatmap (**Fig 2A**).

**Figure 2.**
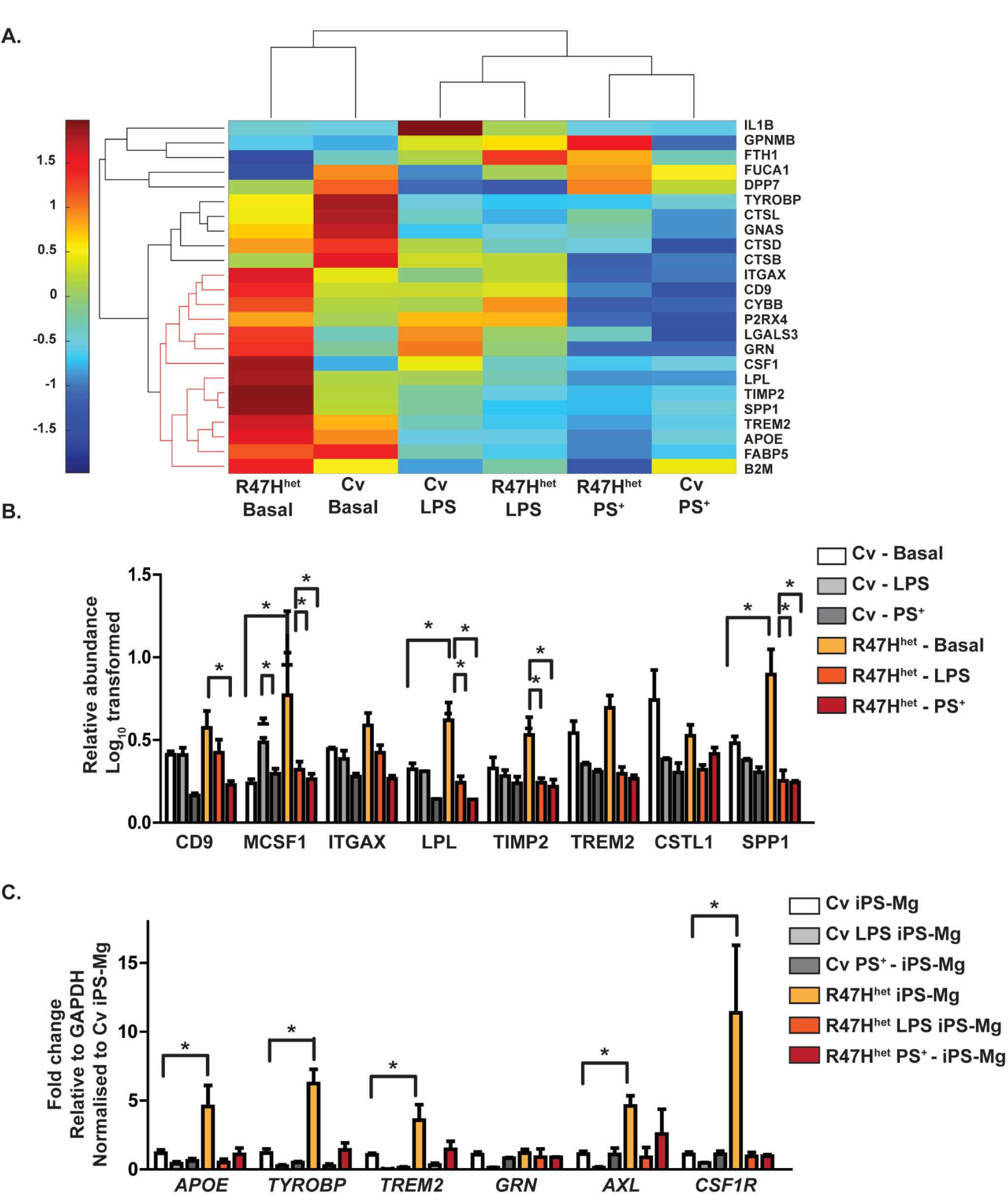
Proteins associated with the DAM microglia signature are found in exosomes. The exosomal proteomic profile was compared with a list of published DAM-related proteins [37–39]. R47H^het^ exosomes contained a subset of DAM-associated proteins (indicated in orange) at higher levels than Cv exosomes (**A**). These proteins were particularly enriched with the TREM2-dependent 2^nd^ stage of DAM [37]. The relative abundance of the DAM proteins in the exosomes was normalised to the abundance observed in Cv exosomes of these 2^nd^ stage DAM-associated proteins (**B**). The expression of DAM associated genes was measured in the iPS-Mg using qPCR to assess whether the changes observed in the exosomes reflect differences at the cellular level or the packaging (**C**). For **A** and **B**, N=3 independent samples analysed through LC-MS/MS for each condition, whilst for **C**, N=4 independent experiments. Two-way ANOVA with * p < 0.05, ** p < 0.01.

A subset of these DAM-associated proteins (indicated in red in the dendrogram) displayed particularly high levels in basal R47H^het^ exosomes (**Fig 2A**). These proteins were linked to the second *TREM2*-dependent stage of DAM microglia [37]. These proteins were plotted individually (**Fig 2B**), confirming the previous trend shown in the heatmap that R47H^het^ exosomes contained higher levels of DAM-related proteins (**Fig 2A**). This suggests that basal exosomes from R47H^het^ iPS-Mg carry higher levels of DAM-like proteins than Cv iPS-Mg. To determine whether this trend was mirrored by the iPS-Mg cells, the expression of DAM-related genes was assessed through qPCR (**Fig 2C**) which indicated that untreated R47H^het^ iPS-Mg showed higher levels of DAM genes, such as *APOE, TYROBP, AXL* and *CSF1R* [37–39]. This suggested that the changes seen in the exosomes are due to changes observed in the cells themselves rather than changes in exosomal packaging.

### Network analysis

So far, we have analysed the levels of exosomal proteins and the similarities following different iPS-Mg treatments. To elucidate more subtle changes in the data, we used the previously published weighted gene co-expression network analysis [31,32], which has also been used for proteomic analyses [40]. This network analysis is based on the hypothesis that proteins whose levels change in a similar way may be functionally related. This allows the network to build different modules, which are made up by proteins that behave similarly and have similar functions, assigning each protein identified to a specific module. The networks were individually generated for exosomes from Cv iPS-Mg and R47H^het^ iPS-Mg (**Supplementary Fig 2A, B** respectively). Fourteen modules, excluding the grey module containing proteins that did not fall into any other modules, were identified in both networks. Out of the identified modules, a few modules, namely the greenyellow and salmon Cv modules and the greenyellow R47H^het^ modules, were shown to be less preserved (**Supplementary Fig 3, 4**), indicating that they may have been influenced by noise in the data. All of the other identified modules appear to be preserved, as indicated through a low *medianRank*, high *Zsummary* score (**Supplementary Fig 3**) and a significant kME correlation between the two different networks (**Supplementary Fig 4**). The correlation between the modules in both networks with the different treatments was investigated to ascertain what functions exosomes could fulfil following stimulation of iPS-Mg (**Fig 3**).

**Figure 3.**
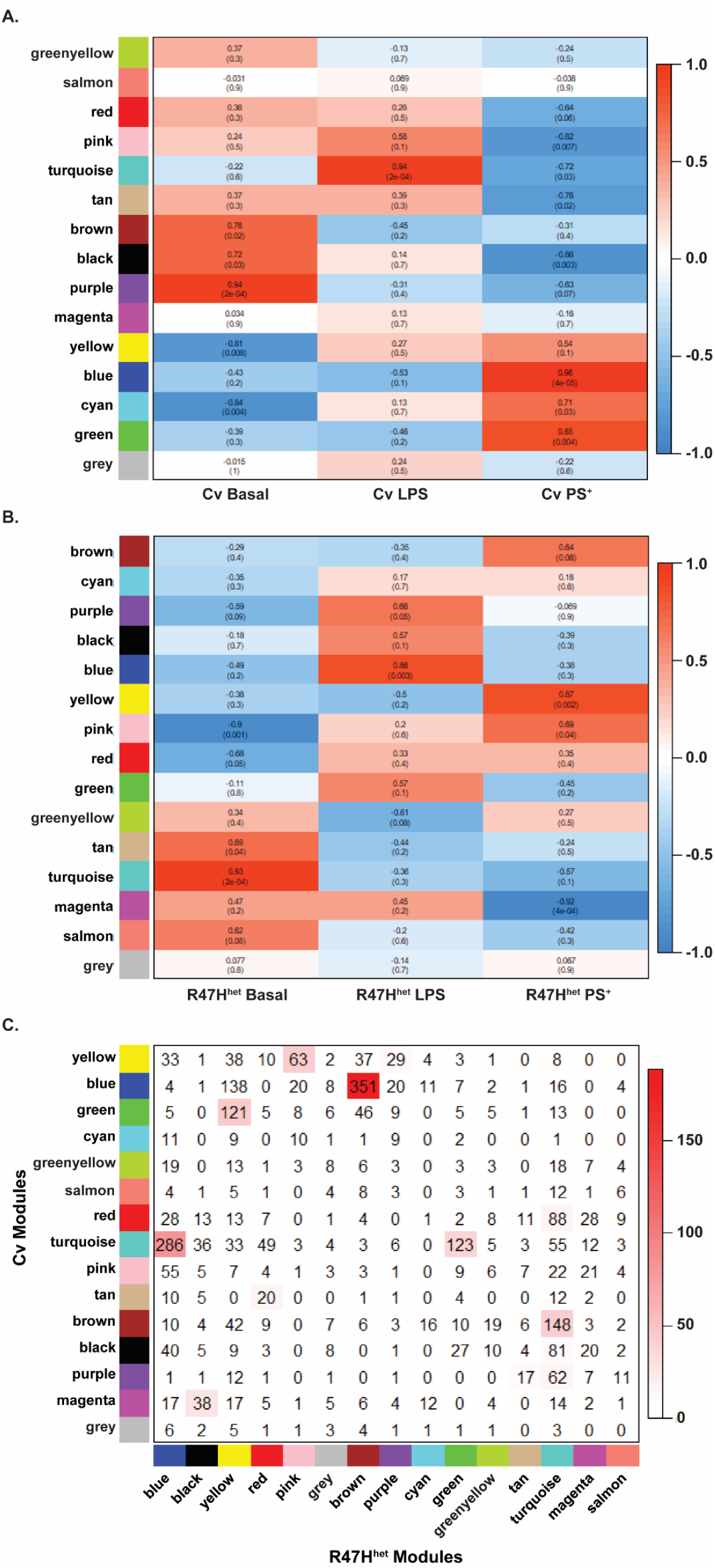
Network analysis. Based on the constructed networks, the relationship between the different iPS-Mg treatments and the identified modules was plotted. The treatment-module relationship was shown for the Cv network (**A**) and R47H^het^ network (**B**). The correlations were indicated in the boxes with the p value in parentheses. The direction of the correlation was colour-coordinated with positive correlations indicated in red and negative correlations in blue. The overlap between the modules identified in the Cv and R47H^het^ network was also plotted (**C**), with the number of shared proteins represented in the boxes and the –log (p value) indicated through the colouring. N=3 independent samples analysed through LC-MS/MS for each condition.

In the Cv network, the modules associated with basal exosomes, brown, black and purple, were enriched for differentiation/development, signalling receptors and the activity of kinases respectively (**Fig 3A**), which was similar to modules which overlapped with the basal conditions from R47H^het^ exosomes, which were enriched for actin organization and differentiation/development (**Fig 3B**). Following LPS treatment of iPS-Mg, exosomes from Cv iPS-Mg were highly correlated with the turquoise module, associated the immune response (**Fig 3A**), whilst the modules associated with the LPS condition in the R47H^het^ network were linked with RNA binding and protein translocation (**Fig 3B**), perhaps indicating different functions of exosomes from Cv and R47H^het^ iPS-Mg. Exosomes from PS^+^-treated Cv iPS-Mg were highly correlated to metabolism, RNA binding and protein translocation, similar to the modules in the R47H^het^ network that were highly correlated to the PS^+^ condition, which were linked to metabolism and proteasome activity.

This analysis suggested both similarities and differences between the modules in the Cv and R47H^het^ networks and therefore, the overlap between the modules identified in the Cv and R47H^het^ networks was plotted (**Fig 3C**). What this shows is that some modules of the two different networks overlap, whilst some modules are exclusive to either network, indicating differences between the two genotypes. Modules associated with the NT conditions in both networks, such as the brown Cv module and the turquoise R47H^het^ module, displayed some overlap suggesting that some functions of basal exosomes appear to be unaffected by the *TREM2* R47H^het^ variant at baseline, albeit with subtle differences. The turquoise immune module in the Cv network is split across two different modules in the R47H^het^ network, again suggesting subtle differences between exosomes from LPS-treated Cv and R47H^het^ iPS-Mg. The modules associated with the PS^+^ treatment display a strong overlap between the two different networks, indicating that there is a strong link to metabolic processes in exosomes from iPS-Mg treated with PS^+^ cells, with potentially some subtle *TREM2* specific differences.

### Microglial exosomes can transmit inflammatory messages

In response to LPS treatment, exosomes from iPS-Mg contained more proteins in the immune cluster identified in the network analysis (**Fig 3**), which is in line with previous studies showing increased inflammatory cytokines in exosomes from LPS-treated microglia-like cells [15]. As inflammatory cytokines can activate other homeostatic microglia, this was further investigated. Exosomes from LPS-treated Cv iPS-Mg significantly increased the expression of *TNF, IL6* and *IL1B* in naïve Cv iPS-Mg (**Fig 4A**), whilst exosomes from PS^+^ treated iPS-Mg had no effect. Exosomes from LPS-treated R47H^het^ iPS-Mg also increased *TNF, IL6* and *IL1B* expression in R47H^het^ iPS-Mg (**Fig 4B**). Interestingly, exosomes from PS^+^-treated R47H^het^ iPS-Mg also increased *TNF* levels in naïve R47H^het^ iPS-Mg (**Fig 4Bii**). Since these effects could be due to carry over of LPS or PS^**+**^ in the exosomes, supernatant (SN) from naïve iPS-Mg was spiked with 100 ng/ml LPS or 2×10^6^ PS^+^ cells before exosomes were extracted. These suspensions therefore would contain any potential carry over without exosomal changes. However, naïve iPS-Mg reacted significantly more to exosomes from activated cells than to spiked exosomes (**Supplementary Fig 5**). This was further supported by endotoxin experiments, which showed that exosomes extracted from basal and LPS-treated iPS-Mg contained negligible levels of endotoxins (**Supplementary Fig 6**). The increase in *TNF, IL6* and *IL1B* expression in iPS-Mg exposed to LPS exosomes (**Fig 4**) is relative to untreated iPS-Mg, however the effect of basal exosomes on inflammatory cytokine expression was also considered. Addition of basal exosomes spiked with LPS or PS^+^ or untreated basal exosomes did not elicit increased levels of *TNF, IL6* or *IL1B* (**Supplementary Fig 5**). Taken together these data support the suggestion that exosomes from LPS treated iPS-Mg were responsible for the increased *TNF, IL6* and *IL1B* expression treated cells (**Fig 4**).

**Figure 4.**
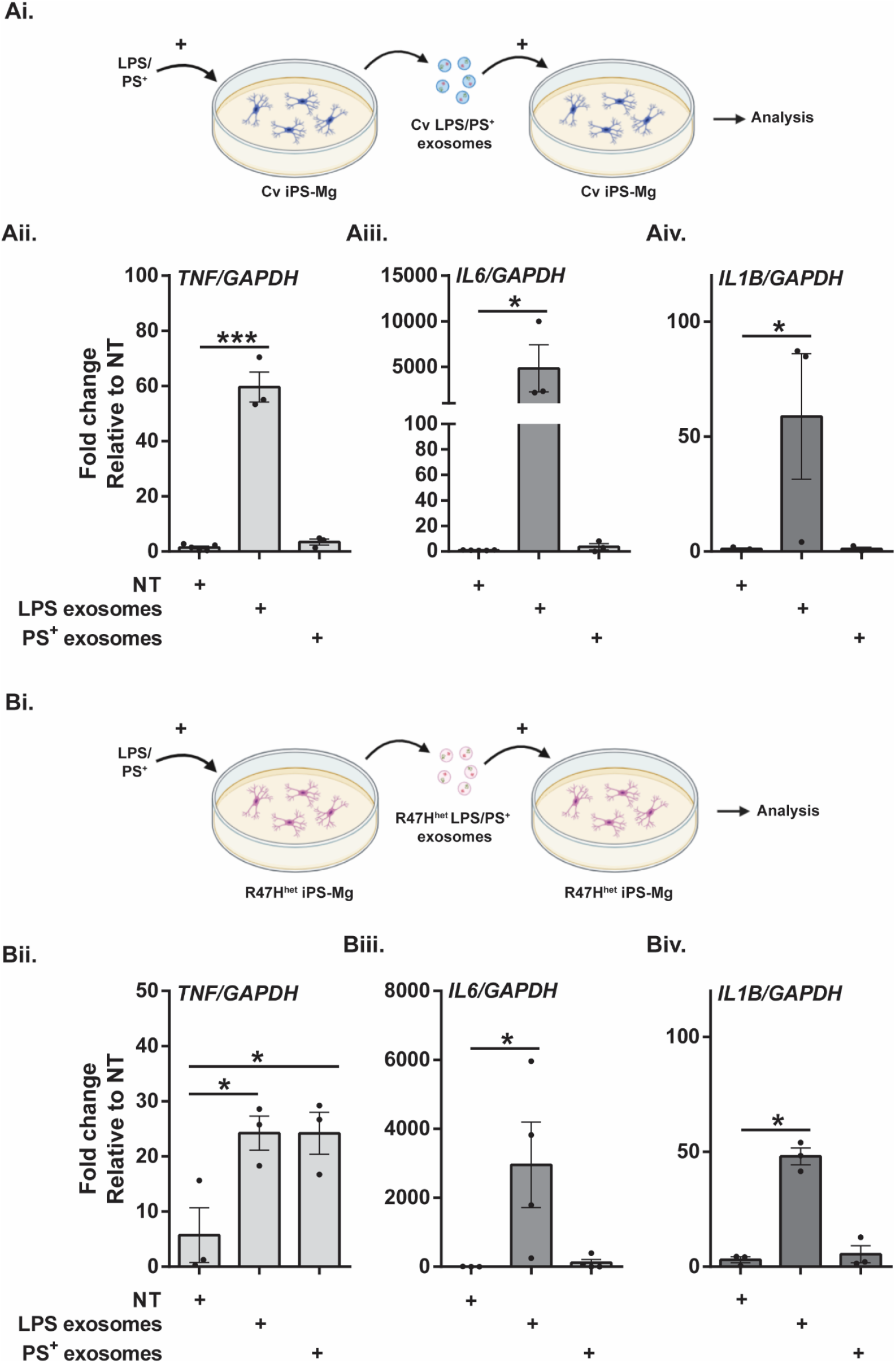
Microglial exosomes can activate naïve microglia. Based on the previous analysis (Figure 3), showing large changes in the immune cluster following activation of iPS-Mg with LPS, the effect of iPS-Mg exosomes on naïve iPS-Mg was measured, specifically, the ability of exosomes from activated iPS-Mg to transmit this information to naïve iPS-Mg. The experimental set-up for **A** and **B** were shown in **Ai** and **Bi** respectively. The expression levels of *TNF, IL6* and *IL1B* were determined by qPCR and were shown for Cv iPS-Mg exposed to Cv exosomes (**Aii, Aiii and Aiv** respectively) and for R47H^het^ iPS-Mg exposed to R47H exosomes (**Bii, Biii and Biv** respectively). N=3 separate experiments with one-way ANOVA. * p < 0.05, *** p < 0.001.

### Basal Cv exosomes can support neuronal development

Because the differentiation and development module was particularly associated with baseline exosomes from Cv and R47H^het^ iPS-Mg (**Fig 3A, B**), and since microglia can support neuronal development [41,42], we investigated the effect of basal exosomes on the development of iPS-neurons, which closely recapitulate human neuronal development [26,43]. When iPS-neurons were exposed to basal Cv exosomes, the length of Tuj1-positive projections was increased significantly, both in comparison with non-treated neurons and those exposed to basal R47H^het^ exosomes (**Fig 5Aii, Aiii**). To control for different numbers of neuronal projections identified in the different ROI, the total length of neuronal processes was normalised to overall Tuj1 staining (**Fig 5Aiii**). In addition, we showed no significant change between both overall Tuj1 staining and voxels identified in the skeleton analysis (**Supplementary Fig 7Ai and Aii** respectively), suggesting that whilst the same number of neuronal processes were imaged in the different conditions, those exposed to basal Cv exosomes formed longer projections.

**Figure 5.**
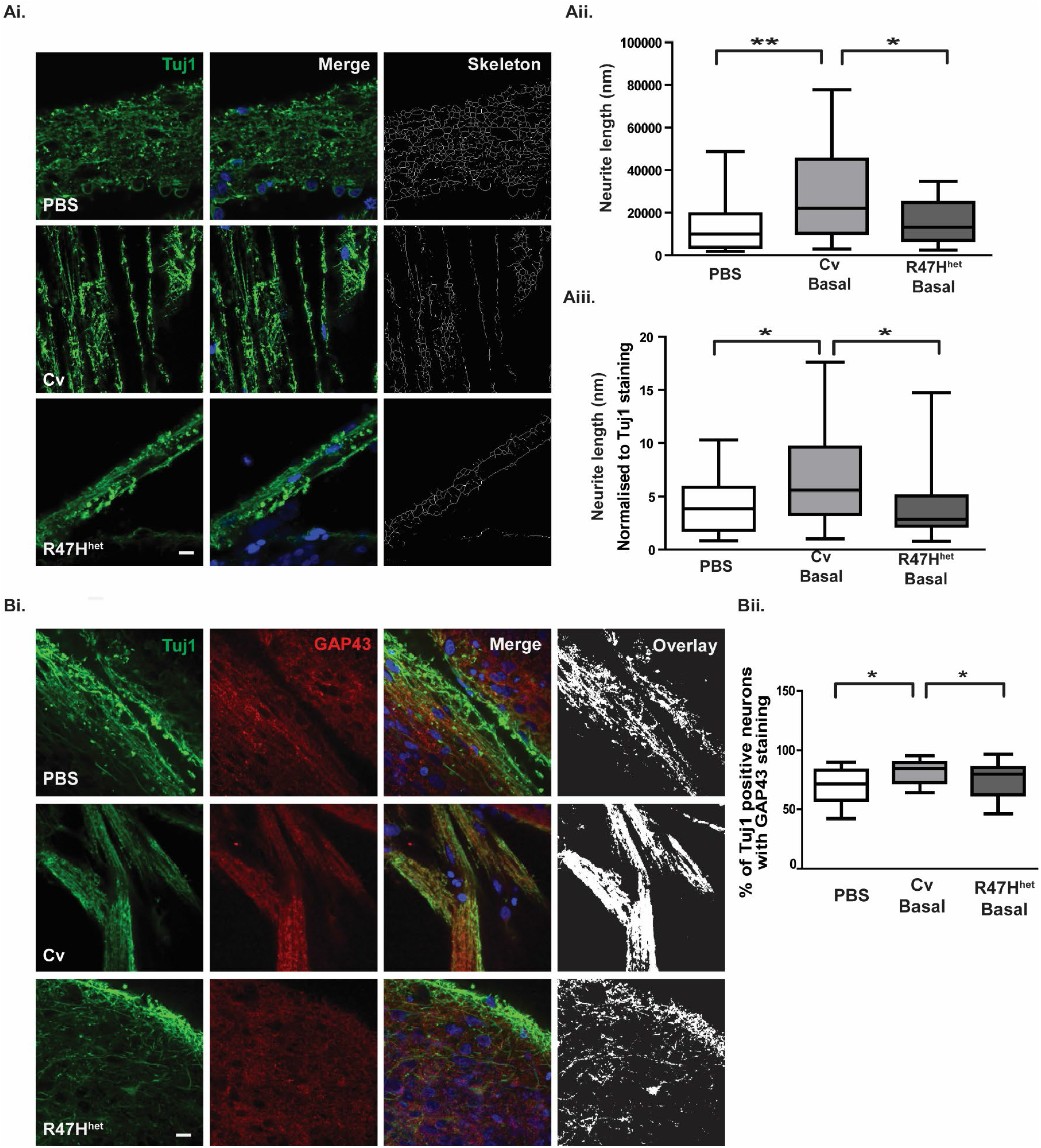
Microglial exosomes support neuronal outgrowth. The ability of exosomes to support neuronal outgrowth was analysed in iPS-neurons exposed to basal Cv and R47H^het^ exosomes. Representative images of iPS-neurons stained with Tuj1 (green) and DAPI (blue nuclear stain) (**Ai**). Plots of skeletonised neuronal process length (**Aii**) and plots of skeletonised neuronal process length normalised to overall Tuj1 staining (**Aiii**) are shown. To verify whether this change was associated with increased neuronal outgrowth, the level of GAP43 staining was quantified in Tuj1-positive projections. Representative images (**Bi**) and following quantification of the overlap) (**Bii**) are shown. N=3 coverslips per condition with 8 ROI analysed per coverslip with one-way ANOVA. * p < 0.05, ** p < 0.01.

One protein that is involved in neuronal process outgrowth, and is found in growth cones is GAP43 [44,45]. To verify that the observed changes in neuronal processes were reflected in increased outgrowth linked to GAP43, GAP43 levels in neurons were quantified. Again, increased levels of GAP43 were observed in iPS-neurons exposed to basal Cv exosomes, compared with non-treated neurons or those treated with basal R47H^het^ exosomes (**Fig 5Bii**).

### Metabolic effects of exosomes on neurons

One of the nodes identified from the analysis of the secreted exosomes involved metabolic functions, in particular in exosomes form iPS-Mg treated with PS^+^ cells (**Fig 3**). To assess whether this function identified through network analysis is indeed translated into biological differences, exosomes were added to differentiated SH-SY5Y, a neuron-like model which display a metabolic profile similar to primary neurons [29]. Basal exosomes from Cv or R47H^het^ iPS-Mg increased the metabolic activity of SH-SY5Y (**Fig 6Aii**). Interestingly, exosomes from Cv iPS-Mg treated with PS^+^ cells induced significantly higher metabolic activity in neurons, than exosomes from R47H^het^ iPS-Mg (**Fig 6Aiii**). Furthermore, changes in ATP levels in SH-SY5Y neurons mirrored the metabolic changes (**Fig 6B**). Basal Cv exosomes increased ATP levels in SH-SY5Y neurons, with a similar trend observed for basal R47H^het^ albeit at lower levels (**Fig 6Bi**). When exosomes were extracted from iPS-Mg treated with either LPS or PS^+^ cells, Cv exosomes evoked significantly higher neuronal ATP levels than their R47H^het^ counterparts (**Fig 6Bii**). These findings were not due to changes in cell death or proliferation of the SH-SY5Y neurons, verified with propidium iodide (PI) (**Supplementary Fig 7Aii**) and crystal violet respectively (**Supplementary Fig 7Ai**).

**Figure 6.**
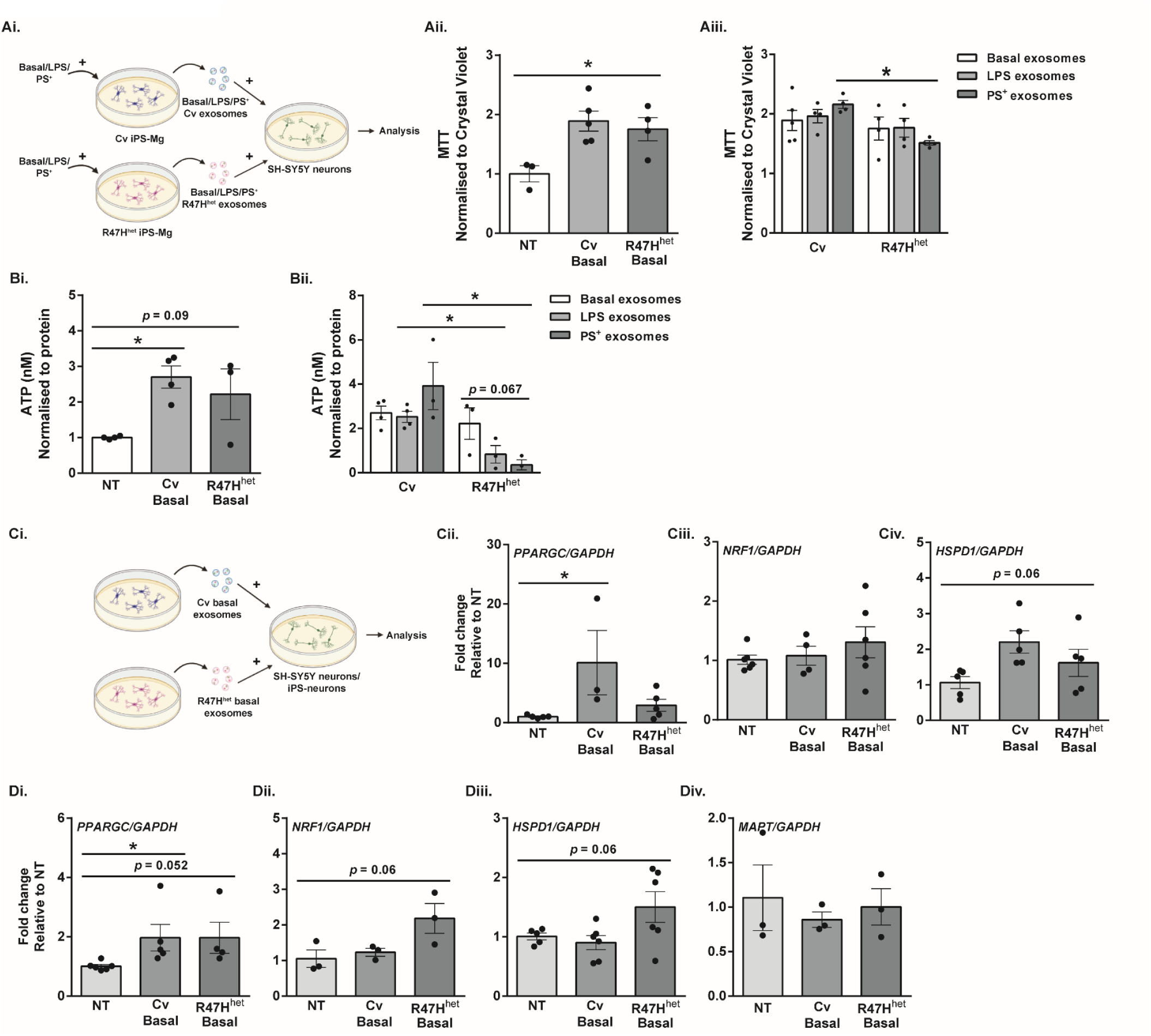
Microglial exosomes can influence neuronal metabolism. Based on the identification of the metabolism cluster and the differences between Cv and R47H^het^ exosomes in this cluster, the effect of exosomes on neuronal metabolism was tested in SH-SY5Y neurons (experimental setup: **Ai**). The metabolic rate was approximated using the MTT assay, (**Aii** and **Aiii**). The production of ATP in the cells, was measured with an ATP assay (**B**) after SH-SY5Y were exposed to exosomes from Cv and R47H^het^ iPS-Mg. Relative expression of various genes in SH-SY5Y neurons and iPS-neurons after they were exposed to basal Cv and basal R47H^het^ exosomes was determined by qPCR, with the experimental setup shown in **Ci**. Expression of *PPARGC, NRF1 and HSPD1* was analysed in SH-SY5Y neurons (**Cii-Civ**) and in iPS-neurons (**Di-Diii**) in addition to the levels of *MAPT* in iPS-neurons (**Div**). N=4 independent experiments for **A** and N=3 independent experiments for **B/C/D** with one-way ANOVA (**Ai, Bi, C, D**) or two-way ANOVA (**Aii, Bii**). * p < 0.05.

To probe how exosomes might influence the mitochondrial function of neuron-like cells, the expression of different genes involved in mitochondrial biogenesis was assessed (**Fig 6C**). Peroxisome proliferator-activated receptor gamma coactivator 1-alpha (PGC-1α) is a key regulator of mitochondrial biogenesis, acting as a transcriptional coactivator for genes involved in mitochondrial biogenesis [46,47]. Through its co-activator the nuclear respiratory factor 1 (NRF1), nuclear genes required for mitochondrial biogenesis can be transcribed [48,49]. Whilst *PPARGC* showed an increased trend in expression following treatment with basal Cv exosomes (**Fig 6Cii**), *NRF1* was not affected by exosome treatment (**Fig 6Ciii**). *HSPD1*, encoding for HSP60 which is involved in the folding of mitochondrial proteins and maintaining ATP production during stress [50–52], showed an increased trend in SH-SY5Y neurons exposed to either basal Cv or R47H^het^ exosomes (**Fig 6Civ**). Due to specific increases in both MTT metabolism and ATP levels following addition of Cv PS^+^ exosomes (**Fig 6Aiii and Fig 6Bii** respectively), *PPARGC, NRF1* and *HSPD1* levels following addition of LPS or PS^+^ exosomes was also tested (**Supplementary Fig 7C**). *PPARGC* and *HSPD1* levels did not further increase in SH-SY5Y exposed to PS^+^ exosomes, but rather the levels fell. (**Supplementary Fig 5C**) indicating that the increase in *PPARGC* levels (**Fig 6Cii**) was specific to basal Cv exosomes.

To verify this in another neuronal model, iPS-neurons were also exposed to basal exosomes from iPS-Mg (**Fig 6D**), particularly as basal exosomes had an effect on the mitochondrial metabolism in SH-SY5Y. *PPARGC* expression increased in iPS-neurons exposed to both basal Cv and R47H^het^ exosomes (**Fig 6Di**) whilst *NRF1* showed slight upregulation following treatment of iPS-neurons with basal R47H^het^ exosomes (**Fig 6Dii**), similar to *HSDP1* (**Fig 6Diii**). Contrarily, *MAPT* showed no changes following exosome treatment (**Fig 6Div**), indicating that overall neuronal number was not affected.

## Discussion

This study investigated how the microglial exosomal proteome was influenced by the stimulus received by microglia as well as by the *TREM2* genotype of the cells. One of the main functions of microglia is to support the differentiation and development of neurons [41,42,53], which they continue to do so throughout lifespan. Exosomes appear to be one pathway through which microglia fulfil these functions, as indicated from our proteomic analysis. In contrast to our previous study where analysis of the proteomic dataset focussed on the top 200 most abundant proteins [22], here we analysed subtle differences in the expression of all 3019 proteins identified by LC/MS using network analysis (**Fig 3**). This specifically implicated that exosomes from homeostatic iPS-Mg could influence development/differentiation, which we further showed here translated into a functional change in iPS-neurons exposed to basal iPS-Mg exosomes including increased outgrowth of neuronal processes, potentially through the increased expression of GAP43. As neuronal process outgrowth is linked to an increased expenditure of energy [54,55], this increase in neuronal process length was mirrored by increased mitochondrial biogenesis and ATP levels in neurons exposed to basal Cv iPS-Mg exosomes. Interestingly, whilst inflammatory cytokines were detected in exosomes of activated iPS-Mg, we did not detect any neurotrophins, in basal exosomes or exosomes from treated iPS-Mg, such as BDNF or nerve growth factor that microglia have been shown to release to support neuronal functioning [56,57]. However, many other microglial factors may be involved in promoting neuronal growth as we showed previously [41], and this is one area of future study.

### Influence of *TREM2* on exosomes from iPS-Mg

In line with our previous study [22], basal exosomes from R47H^het^ iPS-Mg are different to basal Cv exosomes, with basal R47H^het^ exosomes resembling exosomes from LPS-treated iPS-Mg (**Fig 1C**). Others have shown that TREM2 K/O macrophages are more prone to react to low levels of TLR agonists with an exacerbated response [58,59]. Here, the results are more subtle, as one would expect from a heterozygous variant. Thus whilst exosomes from naïve R47H^het^ are already more inflammatory (Fig 1A and C), subsequently exposure to LPS does not significantly change the exosomal profile. This could explain why there was less difference between these two groups when we looked at specific inflammatory markers previously [22].

Whether the changes in exosomal protein content are due to changes in protein packaging or underlying differences in the iPS-Mg themselves was analysed in a subset of identified proteins in this study. In line with another study [60], R47H^het^ iPS-Mg displayed an increased expression of DAM-related genes (overall Fig 2 C is confusing to the rest of the paper) which translated to basal R47H^het^ exosomes containing higher levels of DAM-related proteins, in comparison with basal Cv exosomes (**Fig 2**). Whilst other studies have found an increase in microglia expressing DAM genes in animals treated with LPS [39,61] this was not replicated in this study. This could be due to differences between murine and human microglia, or the time course of the experiments.

### Stimulus specificity include LPS-specific inflammatory changes and PS^+^-specific metabolic changes

Previous studies have shown that exosomal content can change depending on the stimulus the microglia receive [14,15,62]. Here we compared two different stimuli for their ability to influence the exosome proteome. Treatment of iPS-Mg with LPS induced strong associations with the immune clusters, consisting of inflammatory cytokines. This is in line with previous studies [15]. Furthermore, previous research has suggested that cytokines found in exosomes can be active, independent of whether they are found in the lumen or at the exosomal surface [63]. Additionally, exosomes could be a more concentrated delivery system of cytokines to target cells [63], which could elicit a response in either neurons or other microglia exposed to these exosomes. Here exosomes were found to contain inflammatory cytokines which could further activate naïve microglia (**Fig 4**), suggesting a possible pathway of intercellular communication between microglia.

Following treatment of iPS-Mg with PS^+^ cells, metabolic modules were altered. This could be due to PS^+^ cells supplying iPS-Mg carrying *TREM2* variants with additional energy in the form of lipid [22,25]. These changes are also functionally relevant as neurons exposed to exosomes from iPS-Mg treated with PS^+^ cells displayed an increase in metabolism, reflected by increased ATP levels. In particular, the increased expression of PGC-1α and *PPARGC* in neurons exposed to basal Cv exosomes suggests an increase in mitochondrial biogenesis. In addition to this, basal R47H^het^ exosomes also induced upregulation of *NRF1* and *HSPD1*, encoding for HSP60, in iPS-neurons.

### Conclusion/ Outlook

Combining proteomic analysis of exosomal content with downstream functional readouts in recipient cells has allowed us to determine important microglial exosome functions for paracrine signalling pathways, inducing outgrowth of neuronal processes and mitochondrial biogenesis, and autocrine signalling pathways, transmitting inflammatory signals to other microglia. R47H^het^ *TREM2* variant iPS-Mg were shown to secrete exosomes that impair these functions. These findings highlight how dementia risk genes expressed in microglia may translate to changes in the exosomal proteome and point to the possibility of using these as biomarkers.

