## Supplementary Figures for "Differential stimulation of pluripotent stem cell-derived human microglia leads to exosomal proteomic changes affecting neurons"

**Supplementary Figure 1**

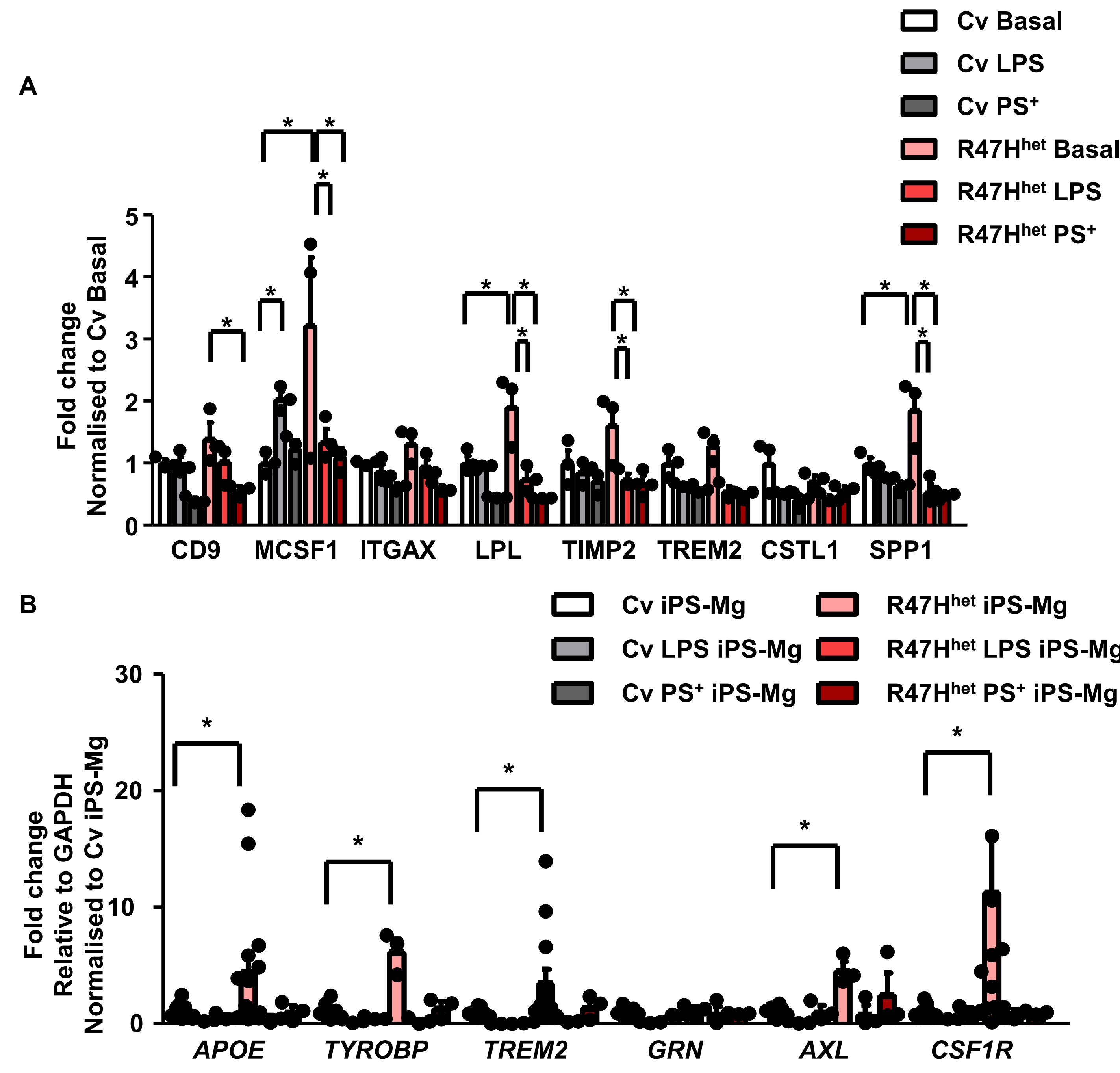


Supplementary Figure 1 – Additional datapoints
Data from Fig 2B and C were replotted in A and B respectively to show individual data points.

**Supplementary Figure 2**

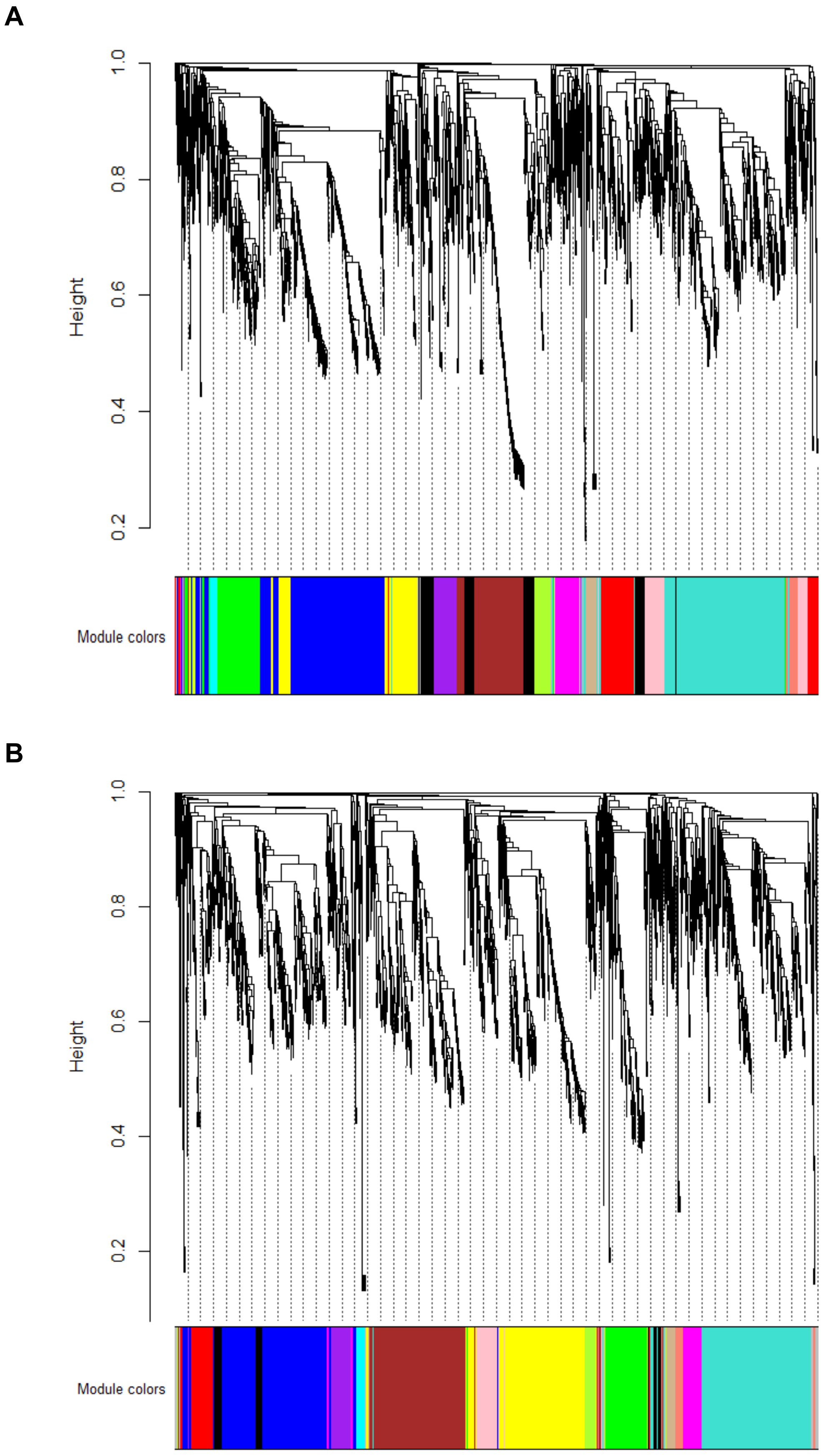


<u>Supplementary Figure 2 – Module building</u>
To generate networks using ProCoNA, proteins were clustered based on their correlation matrix. This was performed for both Cv samples (A) and R47H$^{het}$ samples (B).

**Supplementary Figure 3**

**A**

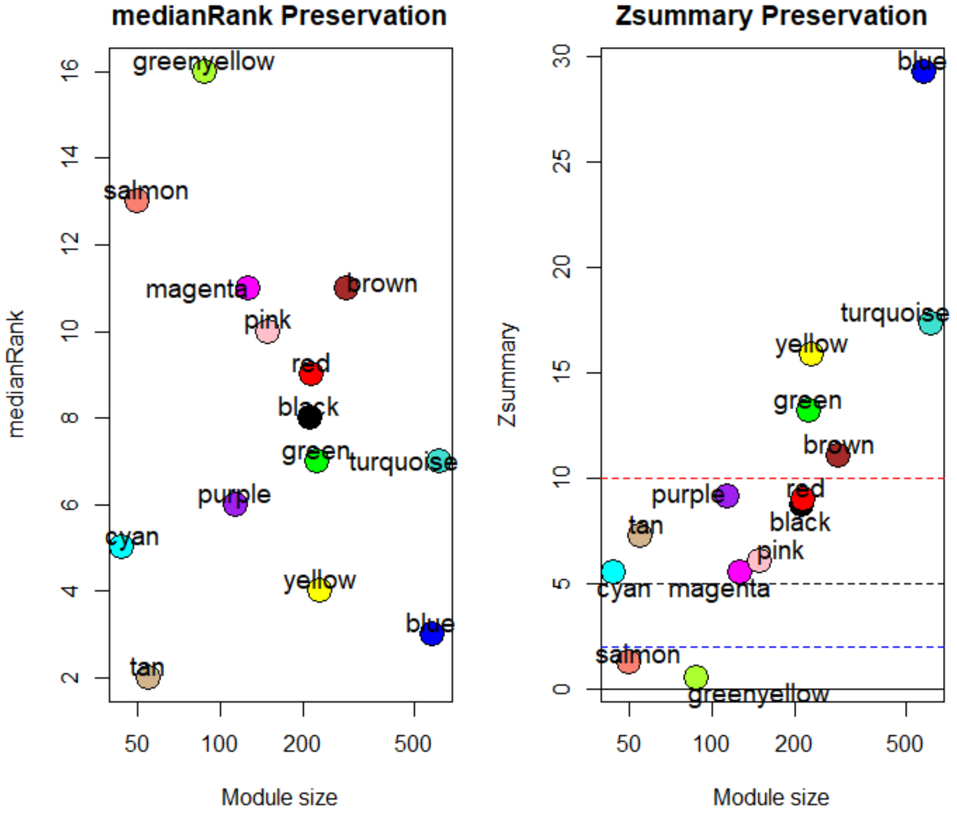


**B**

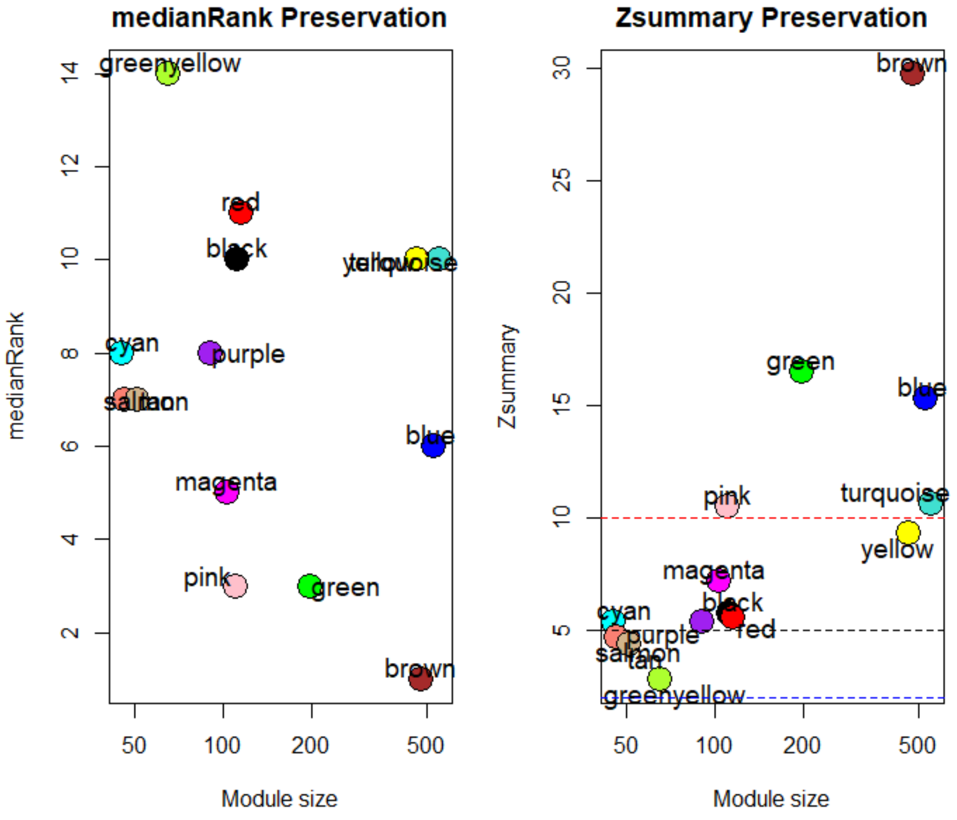


Supplementary Figure 3 – Module preservation

To confirm that the modules identified in the Cv and R47Hhet network are indeed preserved, the medianRank and Zsummary were plotted for each module in the Cv (A) and R47Hhet (B) network.

### Supplementary Figure 4

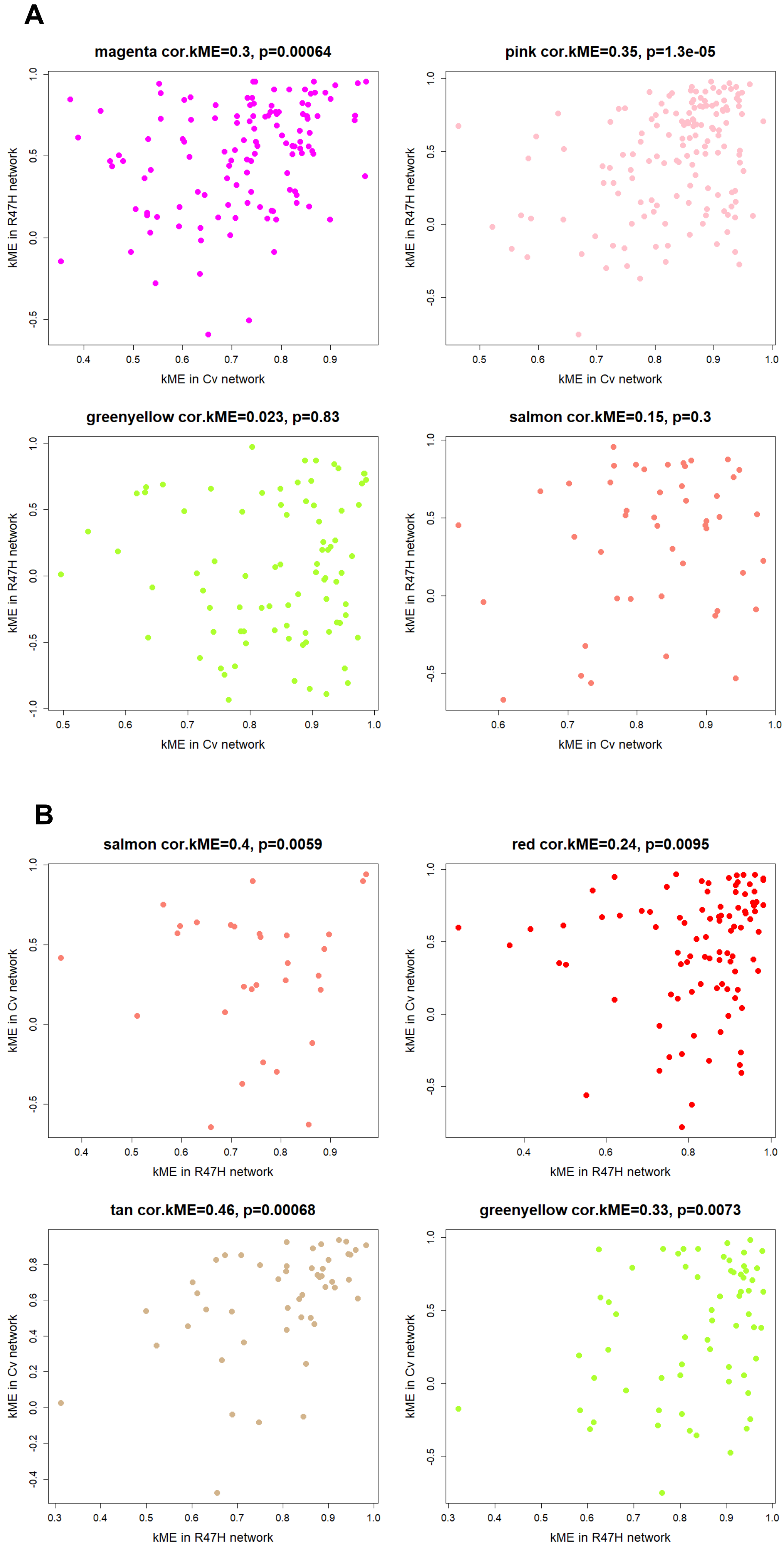


<u>Supplementary Figure 4 – Intramodular correlation</u>

To verify whether the modules identified to only be moderately preserved are indeed preserved in the other network, the kME was plotted for modules in the Cv network (A) and modules in the R47Hhet network (B).

Supplementary Figure 5

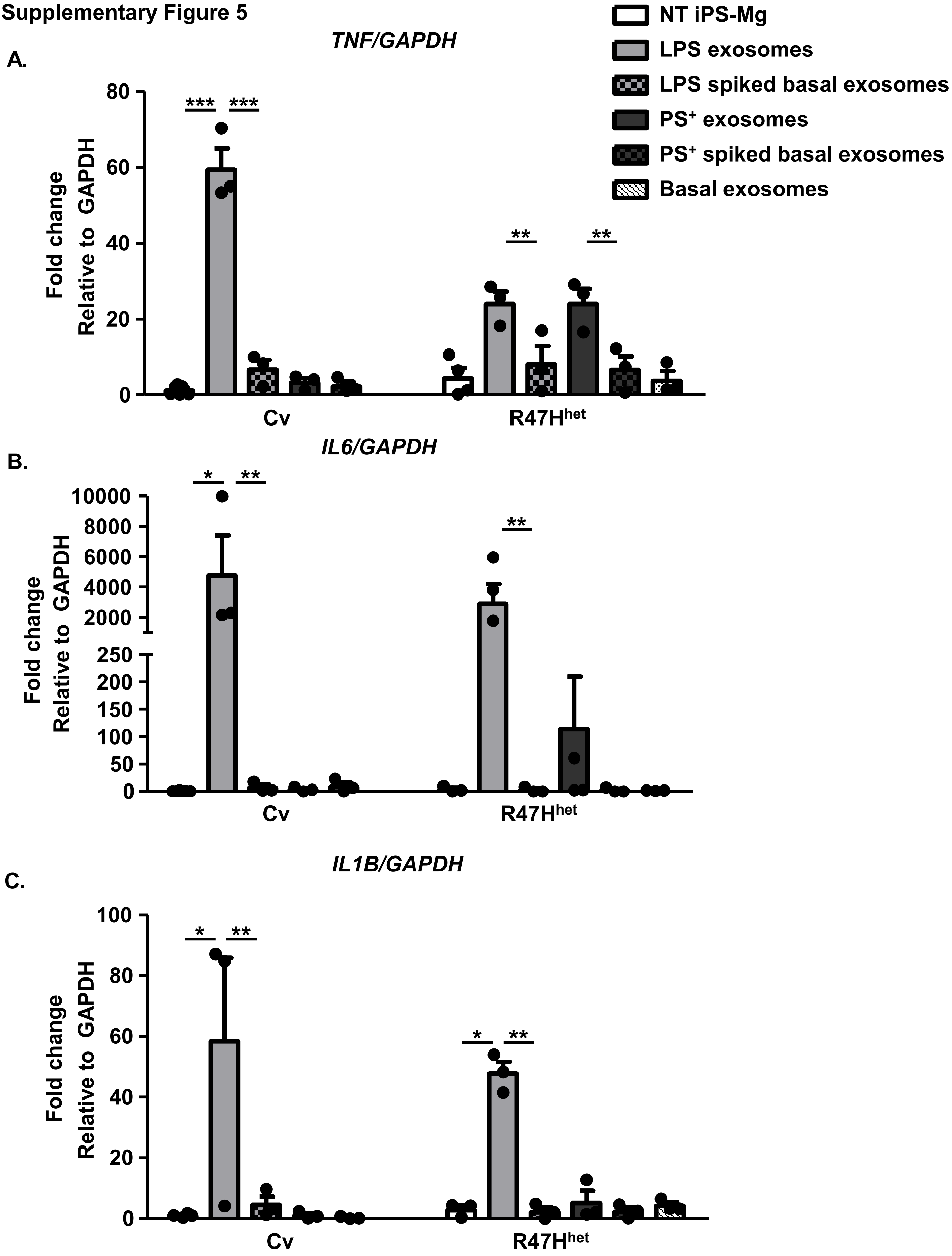


**Supplementary Figure 5 – Effect of spiked exosomes on inflammatory cytokines**

The complete datasets for the expression of TNF, IL6 and IL1B, which were shown in Figure 4, were shown (**A, B and C** respectively). The effect of basal exosomes added to R47Hhet iPS-Mg was also shown. Two-way ANOVA with N=3 and * $p < 0.05$, ** $p < 0.01$, *** $p < 0.001$.

**Supplementary Figure 6**

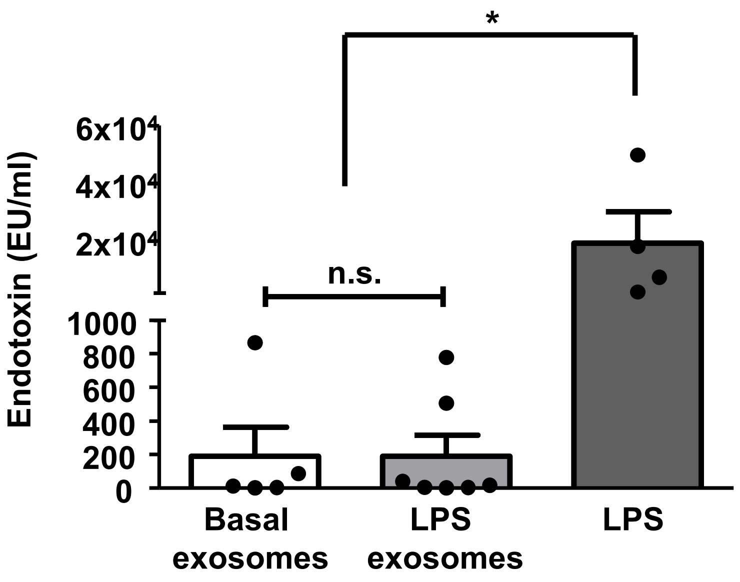


Supplementary Figure 6 – Endotoxin carry-over

Endotoxin levels in exosomes extracted from basal iPS-Mg and iPS-Mg treated with LPS was analysed using an endotoxin test and compared with iPS-Mg medium with 100ng/ml LPS, which acted as a positive control. One-way ANOVA with N=3 and * $p < 0.05$.

### Supplementary Figure 7

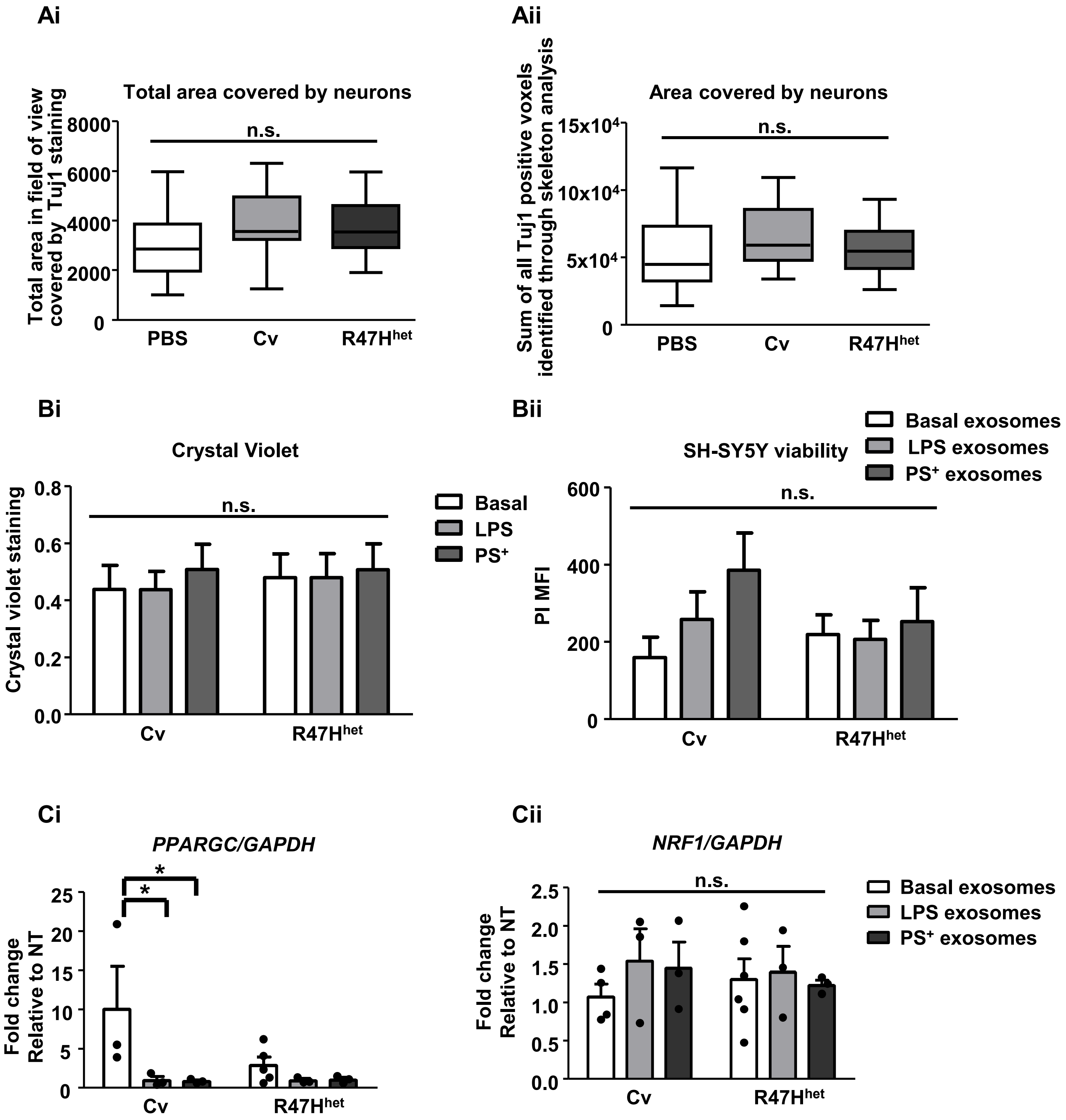


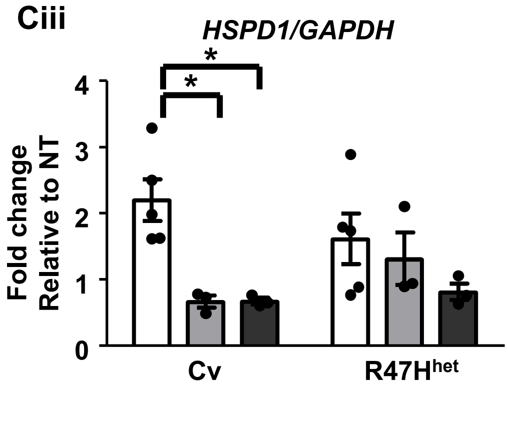


**Supplementary Figure 7**

Controlling for different numbers of neurons present in the image analysis, the total area covered by Tuj1 staining (**Ai**) was plotted in addition to the total number of Tuj1 voxels identified through the skeletonizing plugin (**Aii**). The raw crystal violet data was plotted (**Bi**) to show that overall cell number of SH-SY5Y was not affected when the cells were exposed to exosomes from iPS-Mg. To exclude cell death as another founding variable, SH-SY5Y were stained for PI and MFI analysed using a FACSCalibur (**Bii**). The effect of exosomes from LPS and $PS^+$ treated iPS-Mg on the gene expression of *PPARGC, NRF1* and *HSPD1* in SH-SY5Y was plotted (**Ci-Ciii** respectively. One-way ANOVA (A) and two-way ANOVA (B and C) with N=3 (A, C) or N=4 (B). * $p < 0.05$ and n.s. not significant.
